# Human proton channels accumulate in cholesterol dependent membrane domains via direct interaction with stomatin

**DOI:** 10.1101/2023.09.28.560017

**Authors:** Artem G. Ayuyan, Vladimir V. Cherny, Gustavo Chaves, Boris Musset, Fredric S. Cohen, Thomas E. DeCoursey

## Abstract

Many membrane proteins are modulated by cholesterol. Here we report strong effects of cholesterol depletion and restoration on the human voltage gated proton channel, hH_V_1 in excised patches but negligible effects in whole-cell configuration. Despite the presence of a putative cholesterol binding site, a CARC domain in the human voltage gated proton channel, hH_V_1, mutation of this domain did not affect cholesterol effects. The murine H_V_1 lacks a CARC sequence but displays similar cholesterol effects. These three results all argue against a direct effect of cholesterol on H_V_1. We propose that the data are explainable if H_V_1 preferentially associates with cholesterol-dependent lipid domains, or “rafts.” The rafts would be expected to concentrate in the membrane/glass interface and to be depleted from the electrically-accessible patch membrane. This idea is supported by evidence that H_V_1 channels can diffuse between seal and patch membranes when suction is applied. Suction pulls membrane constituents including H_V_1 into the patch. In whole-cell studies moderate osmotic stretch does not noticeably alter H^+^ currents. Simultaneous truncation of the large intracellular N- and C-termini greatly attenuated the cholesterol effect, but C-truncation only did not. We conclude that the N-terminus is the region of attachment to lipid domains. Searching for abundant raft-associated molecules led to stomatin. Co-immunoprecipitation experiments showed that hH_V_1 binds to stomatin. The stomatin-mediated association of H_V_1 with cholesterol-dependent lipid domains provides a mechanism for cells to direct H_V_1 to subcellular location where it is needed, such as the phagosome in leukocytes.

**Significance:** Many membrane proteins are modulated by cholesterol. Here we explore effects of cholesterol on the human voltage-gated proton channel, hH_V_1. Although we find little evidence for a direct effect, cholesterol was found to exert a strong influence over H^+^ current in excised membrane patches. These effects are explainable by hypothesizing that H_V_1 preferentially associates with cholesterol-dependent membrane lipid domains. We postulate that H_V_1 diffuses within the membrane and is concentrated in such domains that are anchored to the pipette glass by large membrane proteins. We find that H_V_1 co-immunoprecipitates with stomatin, a typical component of cholesterol dependent lipid domains. The association of H_V_1 with lipid domains provides a mechanism for directing H_V_1 to specific subcellular locations to perform specific functions.

## Introduction

Voltage-gated proton channels are ion channels with several unique properties (1); the most critical is strong regulation of their voltage dependent gating by the transmembrane pH gradient (2), resulting in acid extrusion in many cells. They have wide taxonomic distribution, from fungi (3) to insects (4) to humans (5). They play important roles in marine plankton, mediating proton action potentials that trigger the bioluminescent flash in dinoflagellates (6–8). They are crucially involved in the global carbon cycle, where they enable calcification in coccolithophores (9); a role that is threatened by ocean acidification (10, 11). Proton channels perform varied tasks in many human cells, the best established being their synergistic relationship with NADPH oxidase (NOX) that produces reactive oxygen species in phagocytes (12–16), B lymphocytes (17), and other cells.

Recently a significant number of membrane proteins, including ion channels, have been reported to interact directly with cholesterol within the plasma membrane. This interaction may affect the physiologically relevant properties of the protein, as has been found for a number of ion channels (18, 19). In many cases the protein-sterol interaction is facilitated by either a cholesterol recognition amino acid consensus sequence (CRAC) with (L/V)-X_1-5_-(Y)-X_1-5_-(K/R) or the inverse sequence CARC (K/R)-X_1-5_-(Y/F)-X_1-5_-(L/V) domains (20, 21). The human proton channel contains a putative CARC sequence KNNYAAMV (amino-acid residues 131-138) within the extrafacial part of its S2 transmembrane helix and therefore might interact with cholesterol.

## Results

### Phenomenology

#### Cholesterol depletion or addition has profound effects on proton currents in inside-out patches

We explored possible effects of cholesterol depletion from or addition to the membrane using methyl-β-cyclodextrin (MBCD, Methods). When currents were studied in the inside-out patches excised from cells transfected with the human proton channel, hH_V_1, filling the recording chamber with cholesterol-free MBCD solution caused a gradual increase in proton current amplitude on a time scale of tens of minutes (**Fig. 1**). In some cases, the current increased 500% or more upon depletion of cholesterol. The changes in current amplitude were slow and progressive, with the rate of change varying substantially between patches, and did not appear to plateau within the time frame of a typical experiment, making quantitation difficult and unavoidably arbitrary. A further complication was that in most patches, the current amplitude decreased gradually during the first 10-20 min or so after patch excision. We routinely waited until the current appeared reasonably stationary before adding or depleting cholesterol. With these qualifications in mind, depletion of cholesterol increased the current by a factor of 2.79 ± 0.36 (mean ± SEM) in 29 membrane patches over a period ranging 10-40 min. Subsequent replacement of the chamber solution with one containing MBCD saturated with cholesterol caused gradual reduction of the current over a roughly similar time course (**Fig. 1B**). After cholesterol depletion, introduction of cholesterol-saturated MBCD decreased currents by a factor 1.84 ± 0.21 in 14 patches that survived this long procedure. Changing the bath solution yet again to cholesterol-free MBCD caused an increase in current again in 5 cells that survived this long procedure. Despite the large changes in proton current amplitude, the gating kinetics and position of the *g*_H_-*V* relationship remained largely unchanged, as is evident by inspection of **Fig. 1A**. Proton current activation did not change significantly, becoming slightly faster upon cholesterol depletion, with *τ*_act_ decreasing by 13 ± 14% from its initial value in 10 cells in which the kinetics was resolvable.

**Figure 1.**
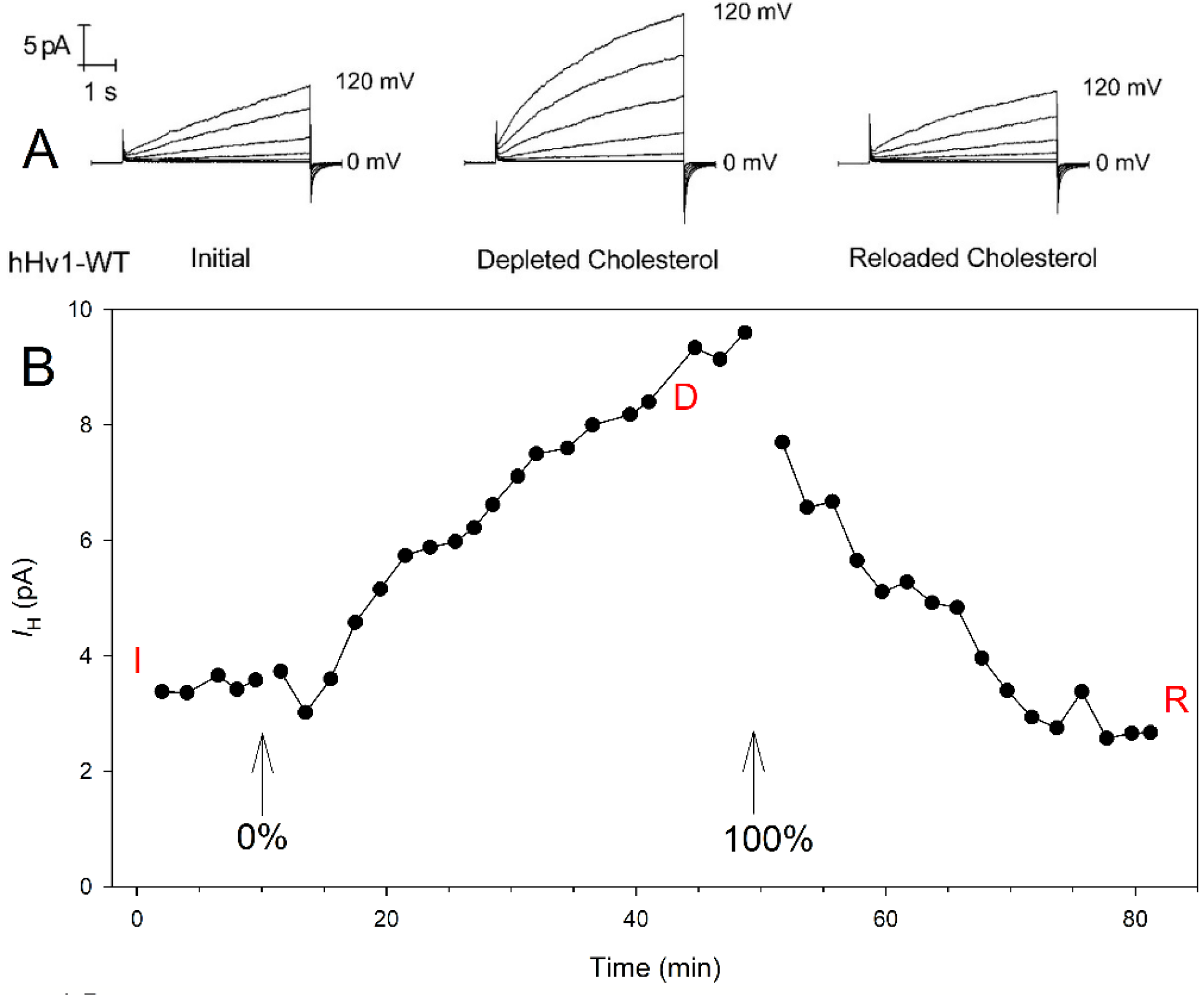
Effects of cholesterol depletion or replenishment on proton currents. (**A**) Families of proton currents recorded in an inside-out patch from a HEK cell transfected with WT hH_V_1. Pulses were applied from a holding potential of −40 mV in 20-mV increments up to +120 mV. (**B**) Maximum current measured during test pulses to +60 mV applied every 30 s in the same patch as **A**. The families in A were recorded at the times indicated in B by “I” (initial), “D” (depleted), and “R” (reloaded).

In 7 cells, cholesterol replenishment slowed *τ*_act_ by 44 ± 21%; neither change was significant (*p* = 0.39 or 0.10). The similarity of proton channel behavior during these large changes in current amplitude strongly suggests that the number of active channels changes, rather than the properties of individual channels.

#### Cholesterol has minimal effects in whole-cell configuration

In stark contrast to the dramatic effects of cholesterol on WT human Hv1 currents in inside-out patch configuration, we found virtually no change in proton current in whole-cell configuration when cholesterol was depleted from the membrane by addition of MBCD to the bath solution. None of the main properties of hH_V_1 changed noticeably. In 6 cells, the maximum current at +40 mV relative to the current in the same cell before depletion, was 0.96 ± 0.06 (*p* = 0.5 by Student’s t-test, hypothesis tested: value change ≠ 1). The time constant of activation of current, *τ*_act_, determined by fitting the rising current to a single exponential, averaged 0.95 ± 0.13 that of the control measurements (*p* = 0.72, *n* = 5). The lack of effect in whole-cell configuration argues against direct modulation of hH_V_1 properties by cholesterol.

A recent report describes an increase in proton current in whole-cell configuration upon depletion of cholesterol by MBCD (22). The reported response was much smaller (∼40%) than that observed here in excised patches, and was not shown to be reversible. We used a several-fold lower concentration of MBCD (∼1.5 mM vs. 5 mM) which may account for the lack of effect on whole-cell currents in the present study. MBCD at concentration of 5 mM or higher is known to extract not only cholesterol but also polar lipids (particularly sphingolipids) from the cell membranes (23). While we cannot rule out possible small effects of cholesterol in whole-cell configuration, the large changes in H^+^ current in excised patches clearly reflect a distinct phenomenon. This phenomenon is the subject of the present study.

#### Increased H^+^ current with cholesterol depletion may reflect increased numbers of channels in the patch

One explanation for the observed effects of cholesterol on proton currents in patches but not in whole-cell configuration is that hH_V_1 molecules diffuse between the exposed patch and the membrane attached to the glass pipette walls. We explored this possibility by applying suction to the pipette, which would be expected to pull more membrane into the conductive part of the patch membrane, increasing the number of channels and hence the current (**Fig. S1A**) (24–26). Given that a previous study reported increased H^+^ current with suction in excised patches (27), it was not surprising that we observed similar increases (**Fig. S1B**). H^+^ current increased during suction, and the increase was larger with greater suction: 10 or 20 in H_2_O (≈2.5 or ≈5 kPa) increased the current reversibly by 33±7 or 60±5 %, respectively (*n* = 4 or 7). Similar effects were explained previously by proposing mechanosensitivity of H_V_1, so that increased membrane tension increased the single channel conductance (27). But the result is also consistent with the idea that the altered currents were due to the addition or removal of electrically detectable membrane area (along with associated channels) from the patch (24–26). The latter interpretation is consistent with the lack of effect of cholesterol modulation in the whole-cell configuration. It is possible that both mechanisms (mechanosensitivity and lateral channel diffusion) coexist.

Proton currents in whole-cell configuration were largely insensitive to membrane stretch. Osmotic swelling activates a ubiquitous chloride current, which is elicited reliably by lowering the external osmolarity by 25% (28) or even by 12% (29). K^+^ currents activated by cell swelling using 17-34% hypoosmotic solutions have also been reported (30). When we reduced the osmolarity by 23% there was little or no change in whole-cell hH_V_1 currents (**Fig. S2**). This was the case whether we kept the ionic strength constant by removing sucrose from the hypotonic solution, or if we doubled the ionic strength, keeping osmolarity constant (**Fig. S2**). Previously, we found minimal effects on proton currents upon reducing the ionic strength by 90% (31). The insensitivity of whole-cell H_V_1 currents to membrane stretch supports the interpretation that a significant fraction of the suction effects in patches (**Fig. S1**) is attributable to diffusion of channels between the seal and the patch membrane.

### Hypothesis

#### Hypothesis: H_V_1 associates with cholesterol-containing lipid domains

The effects of cholesterol depletion or saturation in both hH_V_1 (**Fig. 1**) and mH_V_1 (see below) show clearly that cholesterol influences H_V_1 proton currents, whereas the lack of significant effect in whole-cell configuration indicates that this influence is not direct. To explain these phenomena, we hypothesize that cholesterol affects primarily the number of active proton channels within patches rather than their properties (**Fig. 2**). A significant fraction of the membrane within the excised patch adheres tightly to the glass walls of the pipette and therefore does not contribute to the measured current (24). This adhesion is thought to be facilitated by the interaction of large extracellular domains of membrane proteins (26). A preferential adhesion of these proteins with glass would lead to their accumulation at the glass-membrane interface. Many of these proteins are cell adhesion proteins (*e.g*., CD44, integrins, etc.) which also reside within membrane ‘domains,’ analogous to what have been called lipid rafts (32–35). If proton channels reside preferentially in these domains, they would also become concentrated in the membrane-glass contact area and depleted from the patch. One indication that this is indeed the case was seen immediately after the gigaseal was formed and the patch was excised. The proton current typically decreased several-fold within the first few minutes after patch formation. This can be explained by lateral diffusion of individual membrane domains together with proton channels towards the membrane-glass interface where proteins with large ectodomains will adhere to the glass and thus accumulate preferentially (**Fig. 2A**, Left Side). Eventually a significant fraction of the domains and H_V_1 are depleted from the electrically-conducting part of the patch, reducing the current. “Run-down” of currents after patch excision has been reported frequently for many other ion channels and various explanations have been proposed (36–38), but in some cases it may reflect our proposed mechanism.

**Figure 2.**
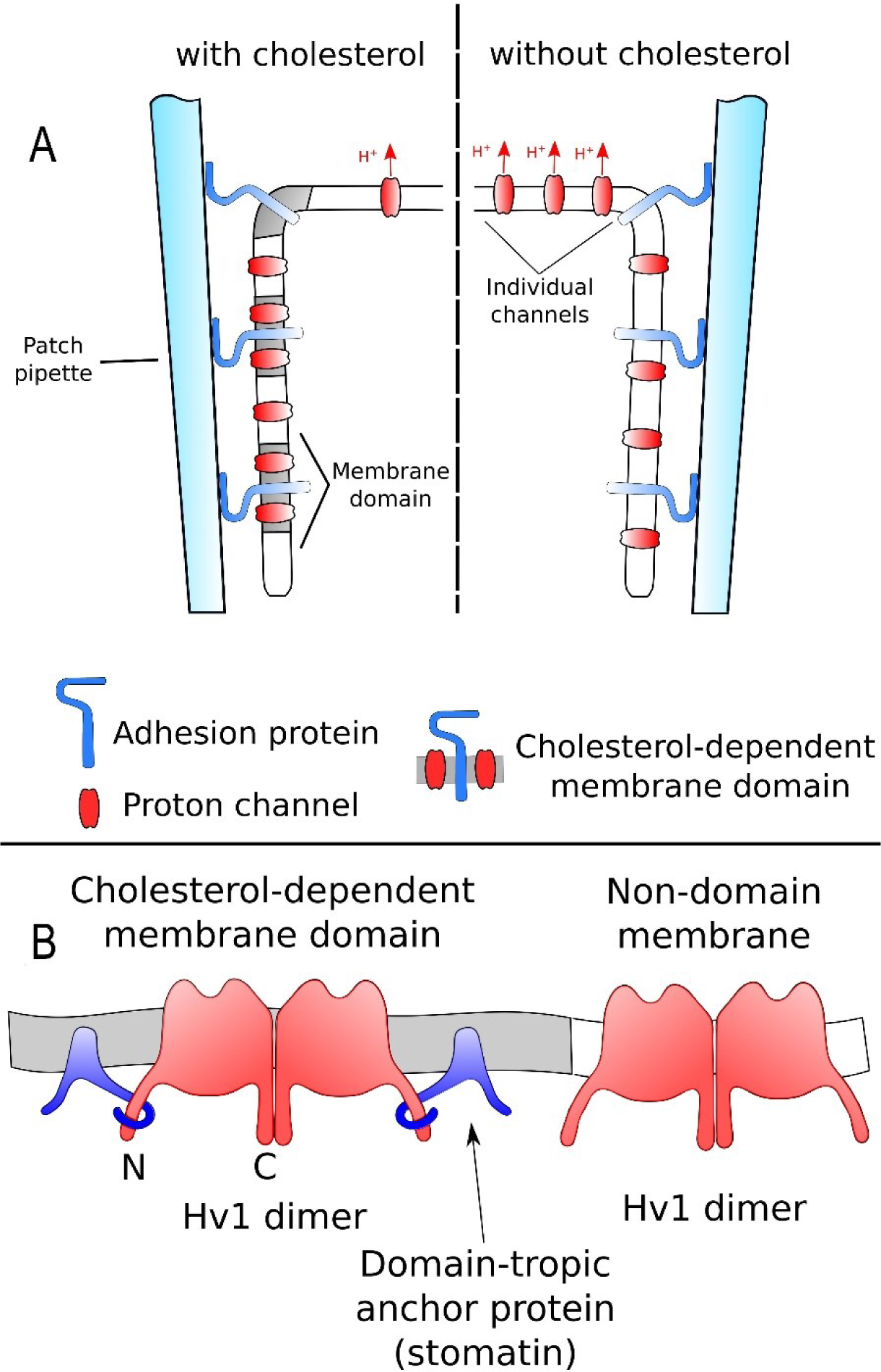
(**A**) Hypothetical explanation for cholesterol effects on proton currents in inside-out patches. (Left) H_V_1 channels are concentrated within membrane domains that also incorporate proteins with large extracellular domains. These extracellular domains adhere to the glass pipette walls when a seal is formed. As a result, a large fraction of H_V_1 channels accumulate at the glass-membrane interface and do not contribute to the proton conductance. (Right) When cholesterol is depleted, membrane domains disperse, releasing the channels to equilibrate through the whole of the patch membrane, increasing the proton current. (**B**) Cartoon illustrating putative binding of H_V_1 to another protein associated with cholesterol-dependent lipid domains.

Some membrane domains (e.g., lipid rafts) are cholesterol-dependent. The depletion of cholesterol from the membrane would cause the dispersal of such domains and liberation of trapped proteins (**Fig. 2A**, Right Side). Proteins with small extracellular motifs, such as hH_V_1 (whose two extracellular linkers are each just 8 amino acids long), would be expected to diffuse laterally and re-equilibrate throughout the membrane similarly to individual lipid molecules (24, 25). For H_V_1 this would increase recorded current as channels move from the seal into the electrically conducting patch membrane.

An independent indication of the lateral mobility of H_V_1 molecules within the patch is the fact that proton current increases reversibly if suction is applied to the patch pipette (**Fig. S1B**). Our results with cholesterol modulation strongly support the mechanism based on the flow of lipids and small proteins, such as H_V_1, within the membrane in agreement with Suchyna and colleagues (26).

### Mechanism of HV1 association with lipid domains

#### The effects of cholesterol are not due to binding to the CARC domain in hH_V_1

It is thought that binding of cholesterol to CRAC/CARC domains controls the partitioning of some proteins to the cholesterol-rich membrane domains (21). In order to test whether cholesterol binding to the CARC domain in hH_V_1 is responsible for the retention of H_V_1 within the membrane domains, we mutated Tyr^134^ of wild-type hH_V_1 to Ala (Y134A). Tyr or Phe residues are essential for cholesterol binding by a CARC domain in other proteins (18, 20). Recordings using inside-out patch configuration demonstrated clearly that this mutant retains its cholesterol sensitivity (**Fig. 3**, yellow bars). The mean change in current amplitude after depletion or replenishment of cholesterol was actually larger than in WT experiments, but not significantly different. Therefore, even if the putative CARC domain in hH_V_1 is functional, it does not control the association of the channel with cholesterol-dependent domains.

**Figure 3.**
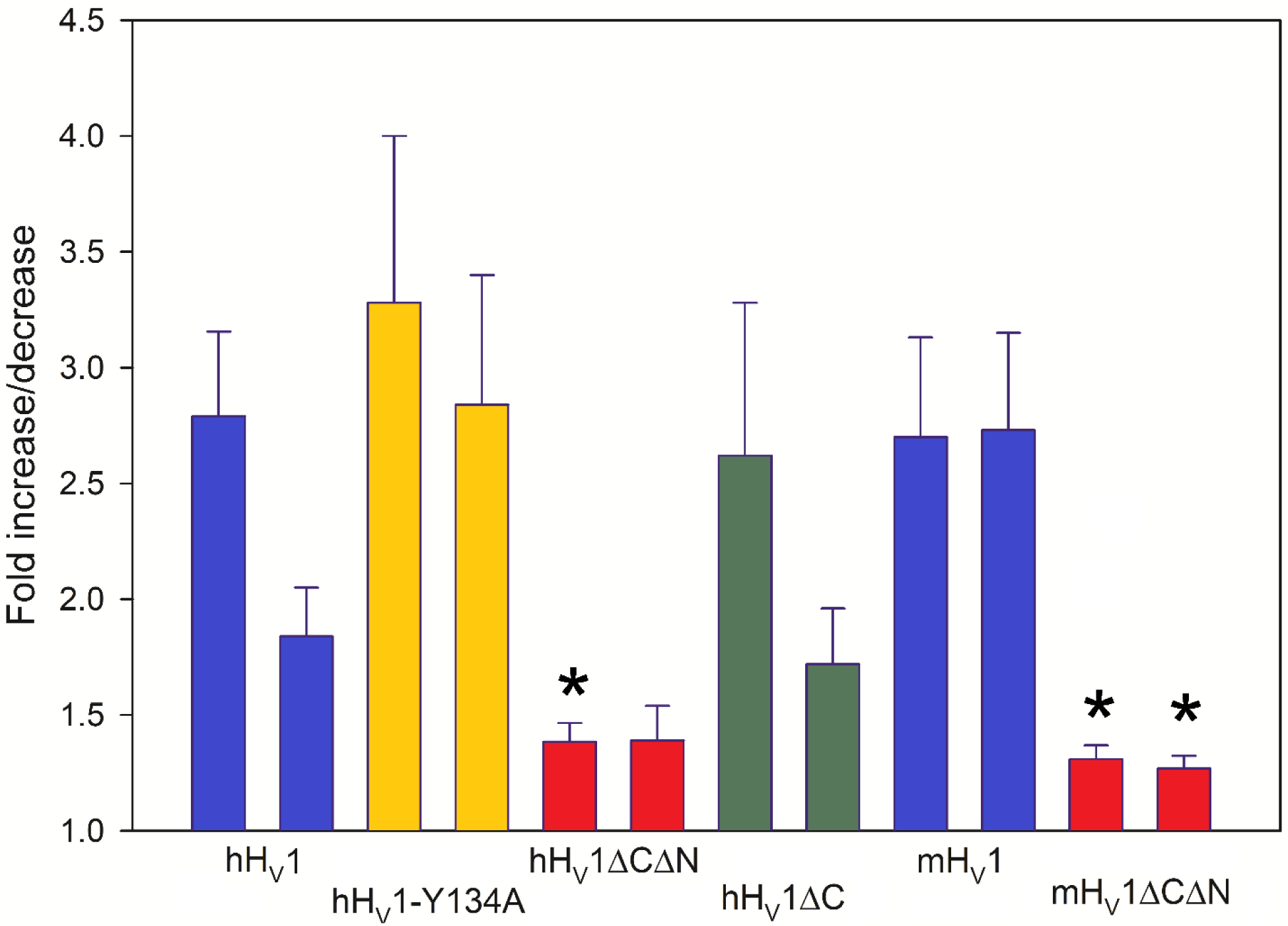
Proton current amplitude during test pulses to +80 mV in the same inside-out patch. Pulses were applied every 30 s from a holding potential of −40 mV. The current measured at the end of each pulse (with the leak current subtracted) is plotted. Arrows show when the cholesterol depleting solution was applied (0%) and later the cholesterol loading solution was introduced (100%). (**C**) Summary of the effects cholesterol (mean plus SEM; n = 29, 14, 10, 4, 12, 9, 9, 7, 5, 4, 7, and 4) in inside-out patches from cells expressing the indicated human or mouse constructs. In each pair, the first column shows the response to depletion, the second replenishment of cholesterol. The Y134A mutation is intended to prevent cholesterol binding to the CARC domain. Starred bars indicate significant difference from the corresponding WT response (by ANOVA, *p* is 0.019, 0.003, and 0.013 respectively).

#### Murine H_V_1 (mH_V_1) exhibits a similar response to cholesterol

Intriguingly, the mouse channel, mH_V_1, while highly homologous with hH_V_1 (78% amino acid sequence identity), lacks a classical CARC sequence at the corresponding site because it lacks K or R at the required position (**Fig. S3**). Nor does any other region in mH_V_1 contain functional CARC or CRAC sequences. It was therefore of interest to see whether cholesterol manipulation still affects the current. **Fig. 3** (mH_V_1, blue bars) illustrates that cholesterol depletion and saturation produced similar responses in WT mH_V_1 channels as were seen in hH_V_1. The fact that the depletion/addition treatment cycle restored mH_V_1 current to its initial value indicates that the treatment did not significantly alter other properties of the patch membrane (*e.g*., phospholipid composition) that could, in principle, also influence H_V_1. The similarity of cholesterol effects in mH_V_1 to those in hH_V_1 argues further against the response being mediated by binding of cholesterol to the CARC domain in hH_V_1.

#### The effects of cholesterol may be mediated by intracellular regions of hH_V_1

In view of the apparent irrelevance of the CARC domain to the association of hH_V_1 with membrane domains, we hypothesized that such association might instead be mediated by direct binding of H_V_1 to another protein that in turn is targeted to cholesterol dependent membrane lipid domains (**Fig. 2B**). This protein-protein interaction is likely to be facilitated by the large cytoplasmic domains (consisting of both the C- and N-termini) of the proton channel. To test this hypothesis, we evaluated whether proton currents conducted by ΔNΔC-truncated mutants (which lack both the intracellular N-terminus and C-terminus) respond to modulation of membrane cholesterol. Both truncated forms of mouse and human H_V_1 exhibited only weak dependence on cholesterol content (**Fig. 3**, red bars) when compared to WT. Therefore, following our hypothesis, one or both termini are essential for the association of H_V_1 with membrane domains.

We then proceeded to test whether an hH_V_1 mutant with the C-terminus truncated (hH_V_1-ΔC) retains the cholesterol sensitivity (**Fig. 3**, green bars). Currents recorded with this mutant indeed responded to the depletion of cholesterol from the patch membrane, as well as its subsequent restoration almost identically to WT, suggesting that the C terminus does not contribute to cholesterol effects. We attempted to conduct analogous studies of an N-terminal truncated mutant, but could not detect viable currents in inside-out patches. Nevertheless, the finding that hH_V_1-ΔC current is modulated by cholesterol but hH_V_1-ΔCΔN current is not strongly implicates the N-terminus of hH_V_1 in the association of the channel with membrane domains. In view of the fact that H_V_1 exists in mammalian cell membranes as a dimer largely due to coiled-coil interactions of the C termini (39–41), it is not surprising that the N terminus would more likely be available to interact with other intracellular protein components (**Fig. 2B**).

#### Identifying binding partners of H_V_1 in lipid domains

Concluding that H_V_1 associates with lipid domains via attachment of its intracellular N terminus to other raft-associated proteins, we wanted to identify its binding partner(s). To narrow down the list of candidates we reasoned that in order to account for the magnitude of the proton current changes observed when cholesterol is depleted, the raft-residing protein that targets H_V_1 to the cholesterol-rich domains must be abundant: H_V_1 has a very low single channel conductance of 20 fS at pH_i_ 7.0 (42), therefore observed changes require a movement of thousands of proton channels and therefore also of raft-targeting proteins bound to them. For example, the increase in current shown on **Fig. 1B** would require a movement of ≈5000 channels from the glass/membrane contact area into the conductive patch. We also reasoned that since proton channels are essential for charge compensation and sustained production of reactive oxygen species (ROS) during phagocytosis (13–16, 43) the raft-tropic protein(s) should also be abundantly present in phagosomes. Flotillins 1 and 2 as well as stomatin fit perfectly into this role: they are all highly abundant in cholesterol-dependent membrane domains and are present in phagosomes (44, 45). To test this hypothesis, we carried out co-immunoprecipitation (co-IP) experiments. Results shown in **Fig. 4** demonstrate that stomatin indeed precipitates when hH_V_1-GFP is used as the bait. We further modified the standard co-IP procedure to characterize the nature of interactions between hH_V_1 and stomatin. A standard co-IP experimental procedure uses a mild detergent to disrupt the cells and solubilize the proteins. Since low temperature (2-4°C) is also used, these conditions are known to result in the formation of detergent resistant membranes (DRMs) - large protein-lipid complexes containing variety of proteins that are held together by interactions that are crucially dependent on cholesterol (32, 46–48). Therefore, the regular co-IP procedure cannot rule out the possibility that hH_V_1 and stomatin are simply present together in the same lipid domain rather than interacting directly or through an adaptor protein. To test for this possibility, we used cholesterol oxidase to convert virtually all cholesterol in the cell lysate into cholestenone (cholest-4-en-3-one) prior to co-IP procedure. This reaction was conducted at 2°C and was run to completion over ∼2 hours. We then repeated the co-IP procedure using this enzymatically treated lysate and found that the results did not change: stomatin co-immunoprecipitates with WT hH_V_1-GFP (**Fig. 4**). Because cholestenone does not support the existence or maintenance of rafts (53, 54), the coprecipitation shows that hH_V_1 associates with stomatin via protein-protein interactions that are independent of cholesterol. In a separate set of patch-clamp experiments, we tested the effects of replacing cholesterol in the patch with cholestenone. To do this the excised patch was exposed to the MBCD solution saturated with cholestenone. Since the solution contained no cholesterol, this caused a rapid removal of cholesterol from the patch membrane and its substitution with cholestenone. This procedure caused the proton currents to increase (1.75 ± 0.17)-fold (*n* = 4, *p* = 0.022), similar to the effect observed when cholesterol is simply depleted (**Fig. 3**). We have also conducted co-IP experiments using the cells co-transfected with hH_V_1-GFP and flotillins 1 or 2 but found no association between these proteins. Together, the results strongly support the role of cholesterol-independent binding of H_V_1 and stomatin in concentrating H_V_1 molecules in lipid domains.

**Figure 4.**
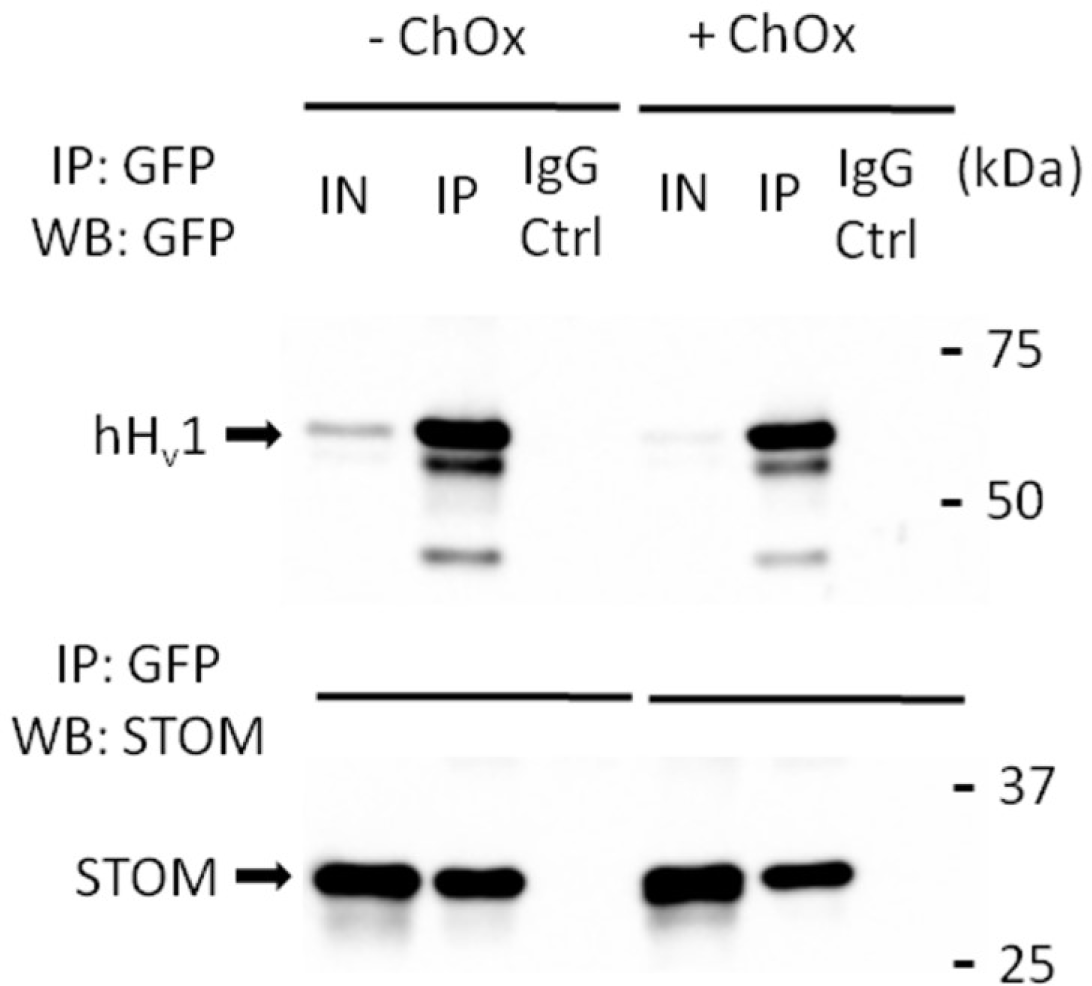
Co-immunoprecipitation of overexpressed hH_V_1-N-GFP and stomatin-N-myc under non-reducing conditions without (-ChOx) and with (+ChOx) cholesterol oxidation. IN: input; IP: immunoprecipitation; IgG: negative rabbit IgG control, WB: Western blot.

## Discussion

### H_V_1 associates with cholesterol-rich lipid domains

Modulation of cholesterol levels had no clear effect on proton currents in whole-cell configuration, but dramatically altered current amplitude in inside-out patches excised from cell membranes. This discrepancy suggests that cholesterol does not directly affect proton currents. Despite the presence of a CARC sequence in hH_V_1 that potentially could bind cholesterol, there was no evidence that the effects of cholesterol that we observed required binding to this site; a mutation designed to disrupt the CARC domain (Y134A in hH_V_1) did not eliminate cholesterol effects. Furthermore, identical cholesterol effects were seen in mH_V_1 which unlike hHv1 lacks a functional CARC or CRAC sequence. Instead, the relationship between the cholesterol content of the membrane and proton current amplitude is interpreted within the framework of a proposed mechanism in which hH_V_1 molecules preferentially associate with cholesterol-dependent domains within the membrane that are anchored to the walls of the pipette.

If H_V_1 preferentially associates with lipid domains as illustrated in **Fig 2**, it would tend to accumulate in the seal region. If cholesterol is then depleted the lipid domains disassemble and release H_V_1 molecules to equilibrate throughout the patch membrane. For this to occur, H_V_1 must be free to diffuse laterally. There is a large literature on the rates of lateral diffusion of membrane proteins, especially ion channels. At just 273 amino acids (31.7 kDa), hH_V_1 is a relatively tiny ion channel, and given its minuscule extracellular loops (both comprise 8 amino acids) it seems very likely that H_V_1 is free to diffuse within the membrane. We support this notion by showing that suction, which would be expected to pull membrane and channels contained therein into the electrically accessible patch, rapidly and reversibly increases proton current. Together, the data strongly support the interpretation that cholesterol affects in excised patches reflect changes in numbers of channels in the electrically exposed membrane.

Evidence consistent with the presence of hH_V_1 within lipid rafts was presented earlier (17). In this work Capasso and colleagues showed that in stimulated B-cells hH_V_1 associates with the B-cell receptor (BCR), which in turn has long been known to reside within cholesterol-dependent ordered membrane domains (lipid rafts) (49). This association does not establish direct interaction between the BCR and proton channel molecules but rather that both proteins are independently targeted to the same ordered lipid environment. Recent work (50) has shown that BCR activation (cross-linking) can induce the formation of relatively large signaling membrane platforms that are able to concentrate or exclude other membrane proteins based on their structure. The reported profile of membrane protein partitioning into these BCR-induced signaling platforms closely follows the one expected for ordered lipid cholesterol-dependent rafts (50). Therefore, it is highly likely that BCR cross-linking causes the accretion of smaller pre-existing domains rather than creating them *de novo*. In case of the association between BCR and hH_V_1 (17) this would mean that hH_V_1 is first targeted by an unknown mechanism to the rafts which then coalesce upon B-cell activation.

Accepting that H_V_1 channel proteins preferentially associate with lipid domains in the membrane, we next ask how this association occurs. The hHv1 protein lacks the obvious features, such as palmitoylation (21), typically associated with partitioning into lipid domains. Direct binding of H_V_1 to cholesterol in these domains also seems unlikely, because cholesterol effects were not related to the presence or absence of a CARC domain. This leaves the possibility that H_V_1 is directed to rafts via interaction with other raft-resident proteins. We next asked which part of the H_V_1 protein was involved in this association. The large intracellular N- or C-termini of H_V_1 seem good candidates, because the extracellular loops are small. The fact that truncation of both N-and C-termini largely eliminated the cholesterol effect in both hH_V_1 and mH_V_1 supports this idea. Truncation of the C-terminus of hH_V_1 alone had no effect. These results indicate that interaction of H_V_1 with rafts is mediated by its N-terminus. Given the coiled-coil interaction of the C termini that enforces dimeric nature of H_V_1 in many species, the N-terminus would be expected to be more accessible to interact with other proteins (**Fig. 2C**).

### Identifying H_V_1 binding partners

We speculated that stomatin and flotillins 1/2, which are well-known components of lipid domains (51), and are required for efficient pathogen elimination via phagocytosis (45, 52), might interact with H_V_1. An anchor protein must be very abundant to bind a significant fraction of H_V_1 molecules in view of the low H_V_1 single channel conductance (42). Stomatin and flotillins 1/2 fulfill these requirements. Therefore, we conducted co-immunoprecipitation studies (**Fig. 4**). Our co-IP experiments clearly show that stomatin co-precipitates with wild type hH_V_1 (GFP-tagged) when a standard co-IP protocol is used. We then questioned whether the observed association of hH_V_1 and stomatin is due to specific protein-protein binding or rather non-specific interaction facilitated by the formation of detergent-resistant membranes (DRMs) (32, 46–48) promoted by the use of mild non-ionic detergents such as Triton X-100, CHAPS, NP-40 for cell lysis as well as low temperatures (to prevent the degradation of the lysate). Immunoprecipitating any of the proteins contained within DRMs would precipitate DRMs as a whole, which would reduce the specificity and significance of the co-IP results. To address this concern, we treated the cell lysate with cholesterol oxidase prior to the co-IP. Cholesterol oxidase retains a significant fraction of its activity at 2°C and converts cholesterol into cholest-4-en-3-one. It has been shown that cholest-4-en-3-one, unlike cholesterol, does not support the formation of liquid ordered lipid phase (53, 54) and so oxidation of nearly all cholesterol eliminates any cholesterol-dependent DRM formation. When we repeated the co-IP experiments after oxidizing cholesterol in the cell lysate, we again found that stomatin precipitates together with hH_V_1 (**Fig. 4**) supporting the hypothesis that cholesterol-independent protein-protein binding determines the association of the two proteins. We found no evidence of association of hH_V_1 and flotillins 1/2 under the same conditions.

Here we report drastic enhancement of proton currents by cholesterol depletion from inside-out patches of membrane containing the human proton channel hH_V_1 that was reversed upon restoration of cholesterol. These effects were observed only in excised patches and thus are indirect. Two results rule out the idea that direct binding of cholesterol to the CARC motif mediates the effects of cholesterol observed here. First, mutation of the CARC region did not alter the effects of changing cholesterol content of the membrane. Second, despite lacking a recognizable CRAC or CARC sequence, the mouse channel mH_V_1 responded to changes in cholesterol identically to the human channel. We conclude that H_V_1 channels preferentially associate with cholesterol-dependent lipid domains. This association requires the intracellular N terminus of hH_V_1 or mH_V_1. Co-immunoprecipitation experiments revealed that stomatin binds to hH_V_1. Although stomatin is known to co-localize with cholesterol-dependent lipid domains and to support the formation of multimeric protein scaffolds (55), and a number of ion channels are thought to also associate with lipid domains, very little evidence exists for direct binding of channels with stomatin. This study provides strong evidence that H_V_1 associates with membrane lipid domains or “rafts” (32) by direct interaction with stomatin, providing a mechanism for the localization of these channels to membrane regions where they are needed, for example in phagosomes or other organelles. Lipid raft microdomains localized in phagosomes contain stomatin and flotillins which are required for normal anti-microbial function (44, 45, 52). Thus, a likely function of stomatin in phagosomes is to recruit H_V_1 channels, which are essential to phagosomal ROS production (13–16, 43).

## Materials and Methods

### Mutation and expression system

HEK-293T cells were transfected with WT, mutated, or truncated human or mouse H_V_1 as described (56). A few experiments with WT channel were conducted using COS-7 cells. No difference compared with HEK-293T was observed. Tyrosine-substituted (Y134A) and N-terminal (1-96 aa) truncated mutants were purchased from GenScript Biotech (Piscataway, NJ, USA). For C- and N-terminal truncations of the mouse channel, mH_V_1, a start codon was introduced at T77 and a stop codon at K217 (V216stop), as described (57), producing mH_V_1ΔCΔN. The C-terminal truncation of hH_V_1 resulted from the T222stop mutation producing hH_V_1ΔC.

### Electrophysiology

Transfected cells were recorded in whole-cell or inside-out configurations at pH 7 at room temperature (∼20–26.5°C) as described previously (56). Bath and pipette solutions were used interchangeably. They contained (in mM) 2 MgCl_2_, 1 EGTA, 80–100 N,N-bis(2-hydroxyethyl)-2-aminoethanesulfonic acid buffer, and 75–120 TMA^+^ CH_3_SO_3_^−^, adjusted to bring the osmolality to ∼300 mOsm, and were titrated using TMAOH. Current records are shown without leak correction. Currents were filtered at 0.5-2.9 kHz using hardware filters and then at 20 Hz using the built-in digital filter of a Heka patch-clamp amplifier (EPC-9 or EPC-10). The maximum proton current was the average value recorded during the last 25 ms of a voltage pulse.

### Changing cholesterol levels

Cholesterol content within the patch membrane was varied using methyl-β-cyclodextrin (MBCD), which has been employed previously to modulate membrane cholesterol (23, 58, 59). Cholesterol was depleted from the patch by filling the recording chamber with a solution of 2 mg/ml MBCD in pH 7 solution. In order to load cholesterol, the same MBCD solution was pre-equilibrated with an excess of crystalline cholesterol and filtered through 0.45 µm syringe filter immediately before filling the chamber. These treatments acutely adjust membrane cholesterol content between two extremes (significantly depleted or overloaded). In order to substitute cholesterol in the patch with cholest-4-en-3-one (cholestenone) the bath solution was replaced with 2 mM MBCD solution saturated with cholest-4-en-3-one (Millipore-Sigma, St. Louis, MO).

### Proton channel co-immunoprecipitation and western blotting

293T cells co-transfected with WT hH_V_1-GFP and STOM-N-myc, or FLOT1-N-myc, or FLOT2-N-myc (Sino Biological, Wayne, PA) were lysed at 2°C in lysis buffer (0.5% CHAPS (Millipore-Sigma), 150 mM NaCl, 60 mM TRIS, 5 mM EDTA, pH 7.4) supplemented with Halt phosphatase and protease inhibitors (ThermoFisher Scientific, Waltham, MA). In some experiments cholesterol was oxidized before the co-IP step in order to achieve this 2.5 U/ml of cholesterol oxidase (CO)(Millipore), 20 U/ml horseradish peroxidase (HRP) (Millipore-Sigma), and 50 µg/ml 10-Acetyl-3,7-dihydroxyphenoxazine (ADHP) (Cayman Chemical, Ann Arbor, MI) were added to the lysate and incubated at 2°C for 3 hours. Inclusion of HRP and ADHP protects the lysate from oxidation by scavenging hydrogen peroxide produced by the CO and allows to monitor the completion of the reaction. Virtually all cholesterol was oxidized in 2 hours. In the experiments using hHv1-GFP as bait, rabbit anti-GFP antibody (cat#50430-2-AP, Proteintech Group Inc, Rosemont, IL) coupled to the Protein-G Dynabeads (ThermoFisher Scientific) was used to capture the protein complexes. Captured complexes were dissociated from the beads in the ×2 Laemmli buffer (Bio-Rad Laboratories, Hercules, CA) and analyzed by SDS-PAGE/western blotting using chemiluminescent substrate (ThermoFisher Scientific) and Azure 300 imager (Azure Biosystems, Dublin, CA). Antibodies used for protein detection were as follows: anti-GFP, anti-stomatin (cat. #678941-1-Ig, Proteintech Group Inc.), anti-flot1, and anti-flot2 (Novus Biologicals, Centennial, Colorado).

## Data, Materials, and Software Availability

All study data are included in the article or <u>SI Appendix</u>. Constructs will be made available.

## Acknowledgments

This work was supported by the R35 GM126902 (TD), Kerscher’sche Stiftung (GC), and R01 GM136777 (FC). The authors appreciate advice on plasmid design from Dan J. Bare. No open access license has been selected.

## Author contributions

Conceptualization of the hypothesis: AA

Experimental strategy and interpretation: AA, FC, TD, BM

Data acquisition, patch-clamp: AA, VC, GC, BM

Data acquisition, immunochemistry: AA

Data analysis: AA, VC, GC, TD, BM

Writing: AA, FC, TD, BM

Approved final manuscript: All authors

## Contact information all authors

Artem G. Ayuyan

Vladimir V. Cherny

Gustavo Chaves

Boris Musset

Fredric S. Cohen

Thomas E. DeCoursey

## Supplementary Material

**Figure S1.**
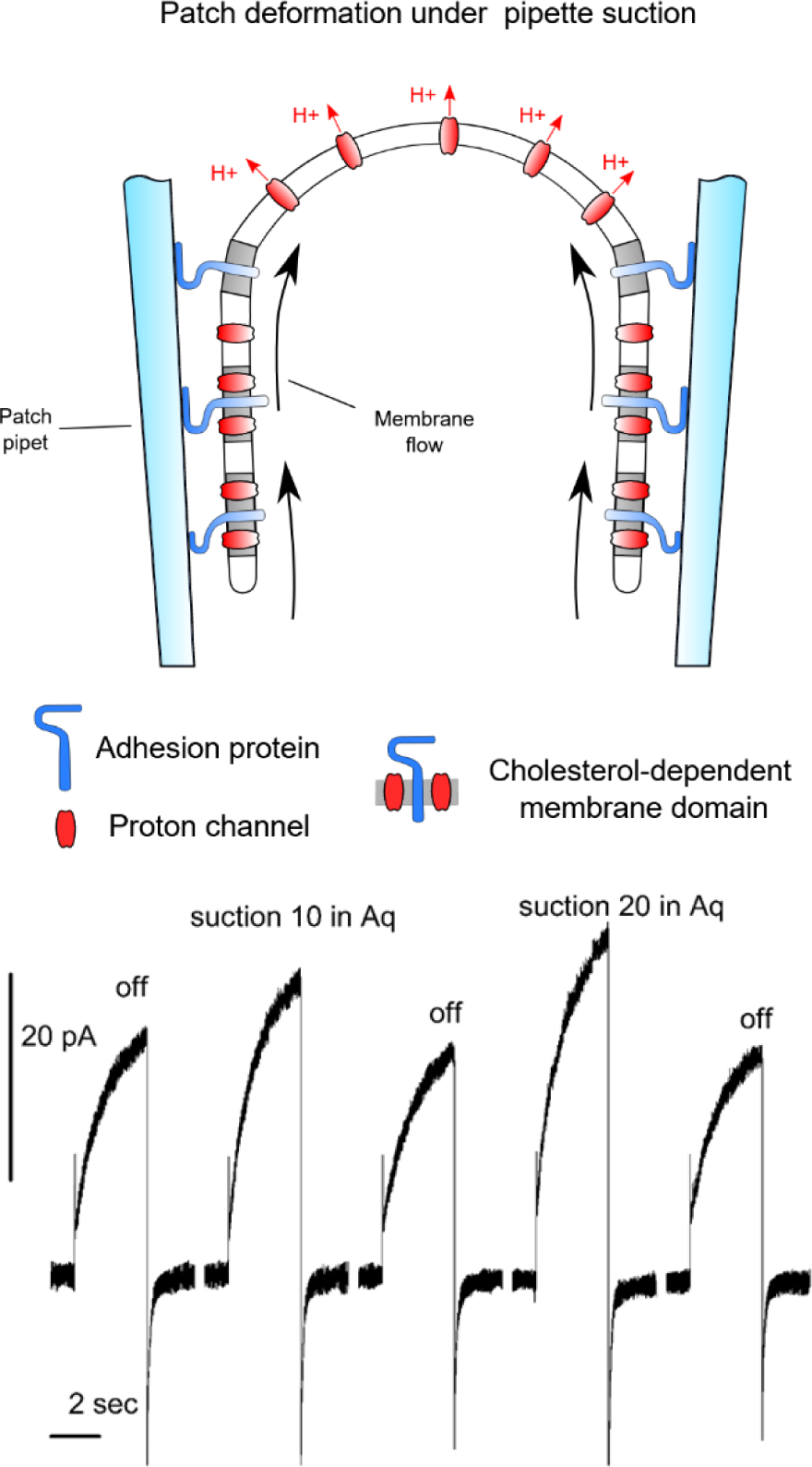
(**A**) Suction applied to the patch would increase the surface area of exposed patch, increasing the proton current in direct proportion to the increased membrane area. Patch area increased due to diffusion of lipids and small proteins, including a fraction of H_V_1 channels not bound within the membrane domains. (**B**) Membrane stretch increases hH_V_1 proton current in excised patches at pH 7//7. Test pulses to +60 mV were applied to an inside-out patch from *V*_hold_ = −40 mV and the indicated suction (in inches of water) applied to the pipette interior.

**Figure S2.**
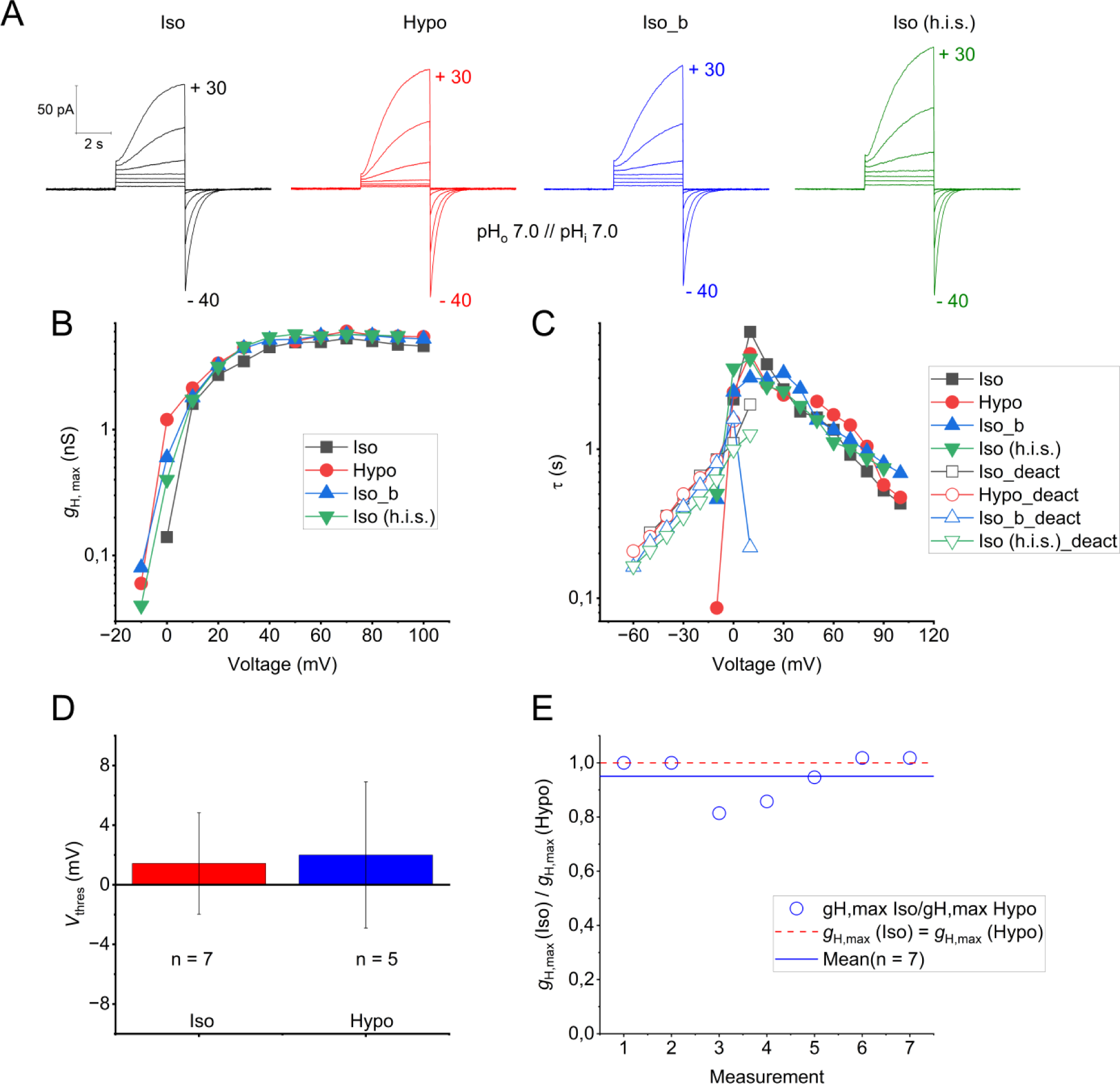
Osmotic effect on proton currents in whole-cell patch clamp configuration. A) Whole-cell patch clamp measurement in the same cell, at hypotonic (231 mOsM, Hypo), isotonic (300 mOsM, Iso), and isotonic with 25% higher ionic strength (300 mOsM, Iso (h.i.s)) conditions. The pH of the cytosolic and the extracellular solution was 7.0. Back buffer exchange to control isotonic solution (300 mOsM) is labelled as Iso_b. The progression of the measurement is shown from left to right. The total duration of the experiment was 1 hour (video recording available!). B) The maximal proton conductance obtained by different pulse lengths (from 1 s to 8 s) of the experiment shown in **A** is plotted against voltage. C) Time constants of activation (solid symbols) and deactivation (open symbols) of the experiment depicted in **A**. D) Comparison between the mean values (± S.E.M.) of the threshold of activation (*V*_thres_) of proton currents under isotonic (300 mOsM, Iso) and hypotonic (231 mOsM, Hypo) conditions. Under isotonic environments, the proton currents activate at 1.4 ± 3.4 mV (*n* = 7) while at hypotonic conditions activate at 2.0 ± 4.9 mV (*n* = 5). All experiments were done at symmetrical pH_o_ = pH_i_ = 7.0. E) To analyse if osmolarity has an effect on the maximal proton conductance (*g*_H,max_) at the steady-state, ratios of isotonic, *g*_H,max_ (Iso) to hypotonic, *g*_H,max_ (Hypo), at the same voltage are plotted. Each point represents a single experiment, the dashed line represents no effect (*g*_H,max_ (Iso) = *g*_H,max_ (Hypo)). The blue line shows the mean value (± S.E.M.) of 0.95 ± 0,03 for the total of experiments (*n* = 7). All experiments were done at symmetrical pH_o_ = pH_i_ = 7.0.

### Solutions composition

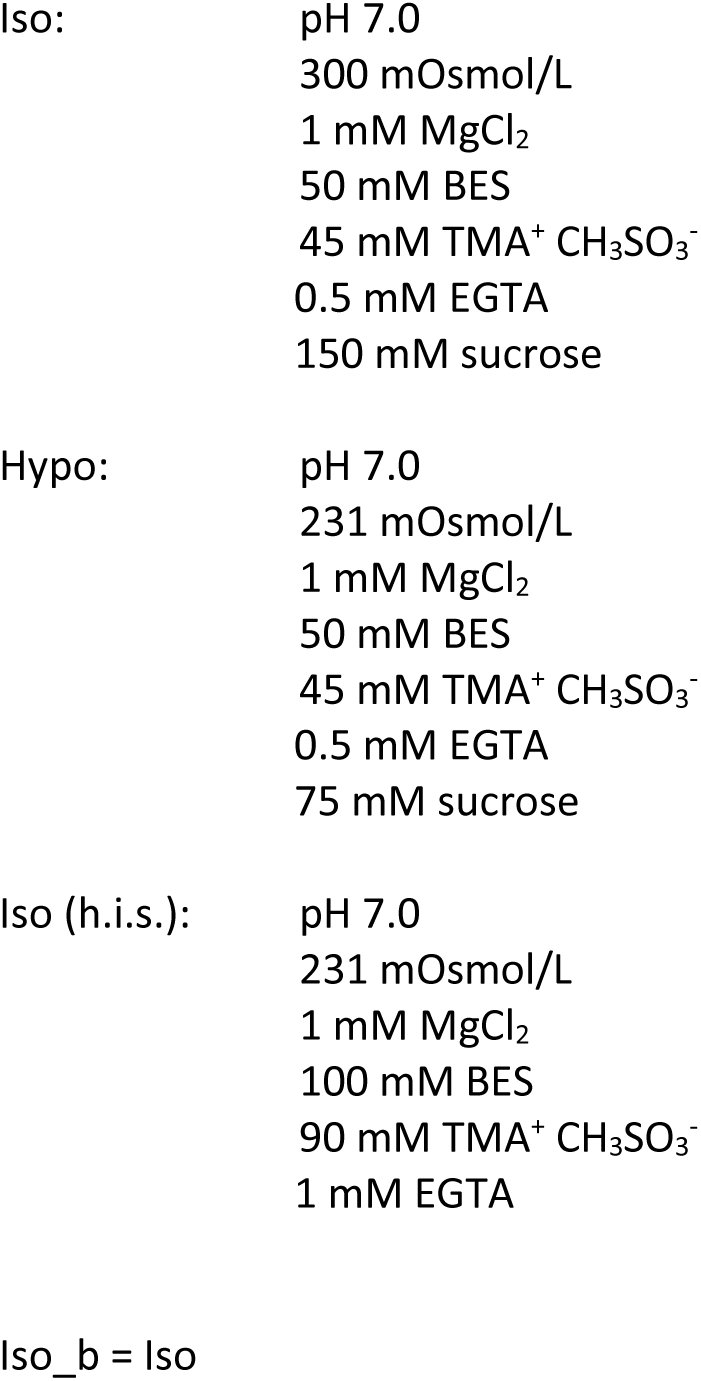

**Figure S3.**
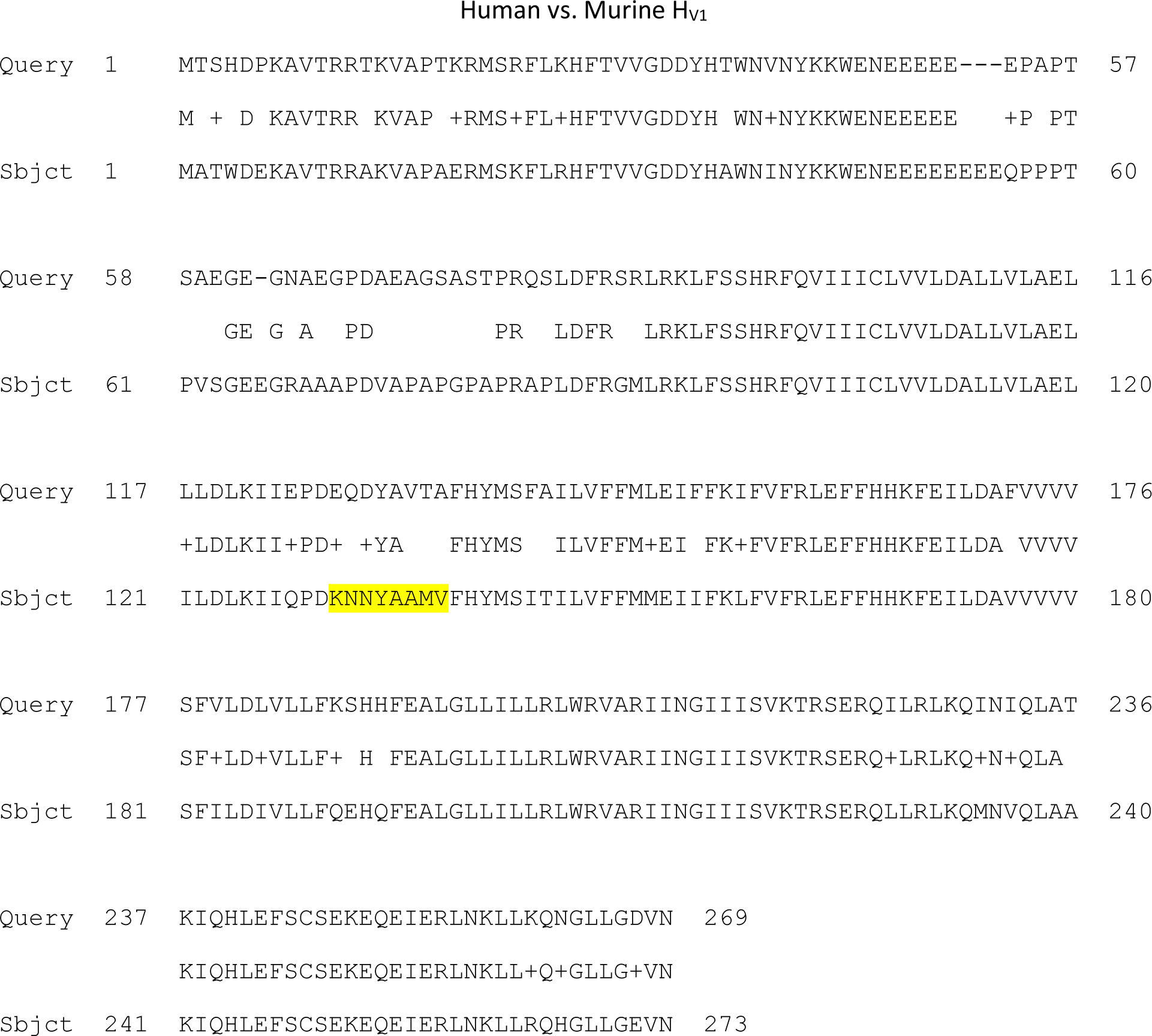
Alignment of mouse “Query” and human “Subject” amino acid sequences. Yellow highlighted region shows a CARC sequence is present in hH_V_1 but not mH_V_1. There are CARC-like sequences in the N terminus of both species, but because they are likely not within the membrane, they would be non-functional as cholesterol binding sites.

**Figure S4.**
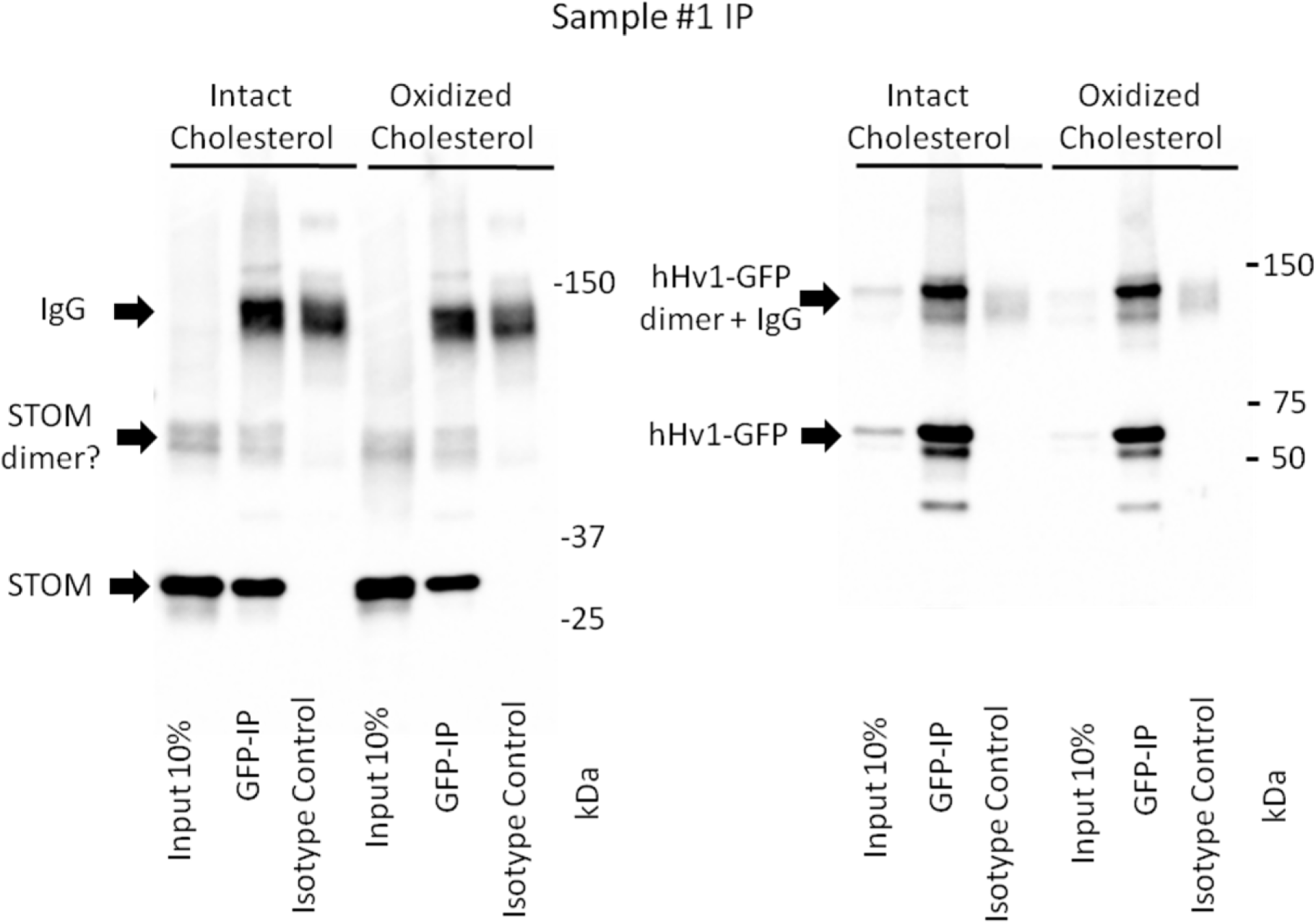
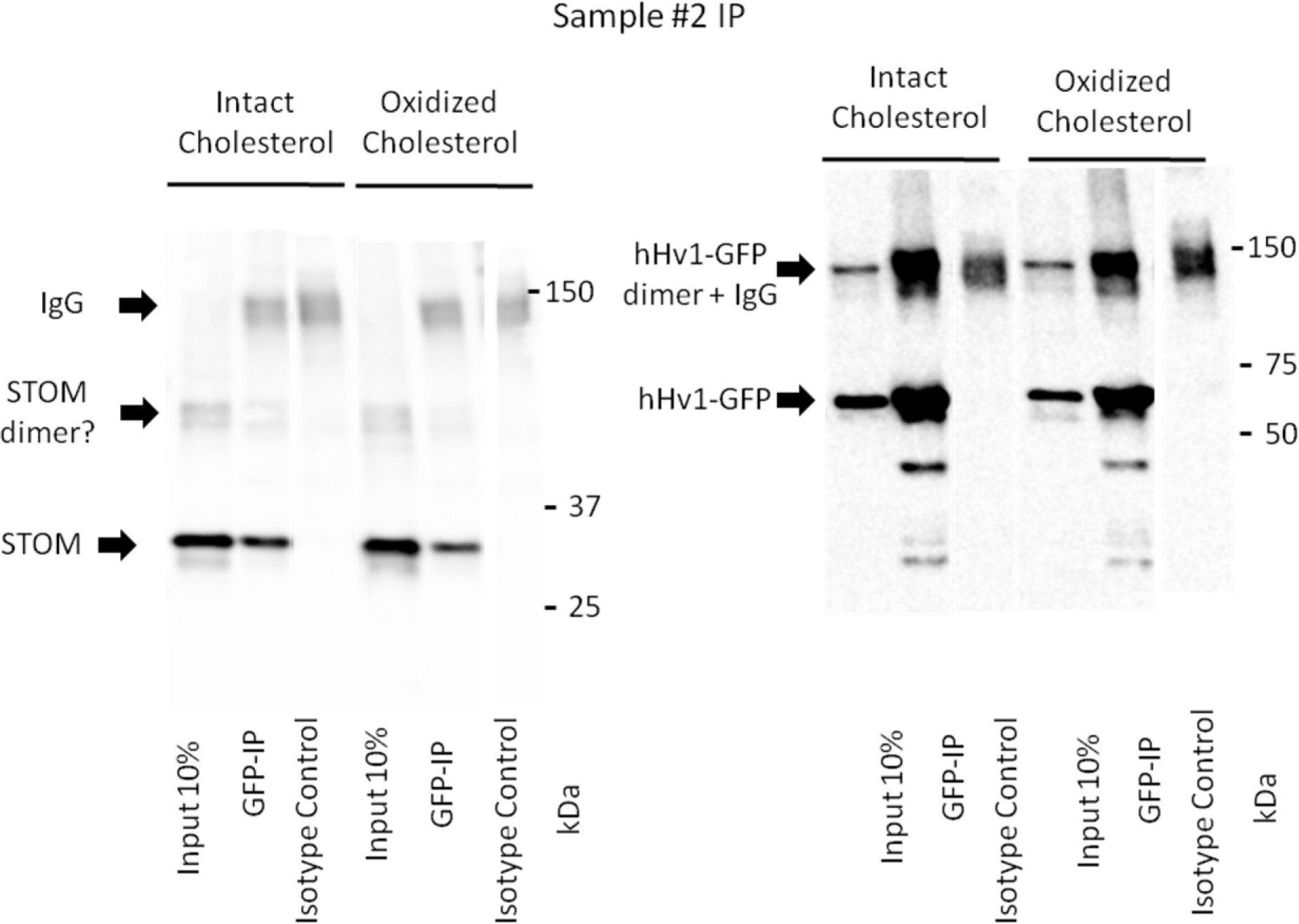
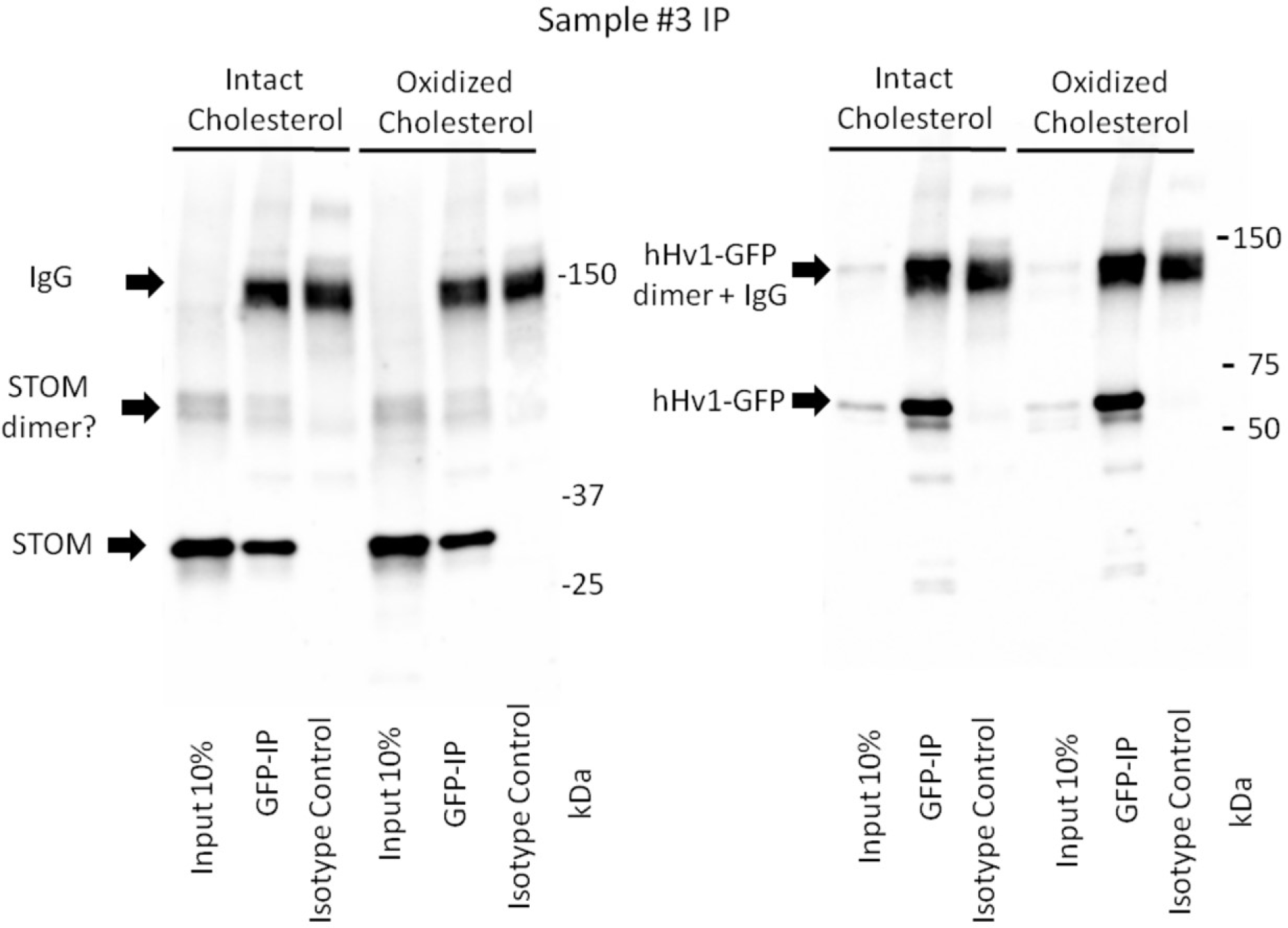
Full, uncut, IP gels.

